# Biophysical modeling of the SARS-CoV-2 viral cycle reveals ideal antiviral targets

**DOI:** 10.1101/2020.05.22.111237

**Authors:** Brian T. Castle, Carissa Dock, Mahya Hemmat, Susan Kline, Christopher Tignanelli, Radha Rajasingham, David Masopust, Paolo Provenzano, Ryan Langlois, Timothy Schacker, Ashley Haase, David J. Odde

## Abstract

Effective therapies for COVID-19 are urgently needed. Presently there are more than 800 COVID-19 clinical trials globally, many with drug combinations, resulting in an empirical process with an enormous number of possible combinations. To identify the most promising potential therapies, we developed a biophysical model for the SARS-CoV-2 viral cycle and performed a sensitivity analysis for individual model parameters and all possible pairwise parameter changes (16^2^ = 256 possibilities). We found that model-predicted virion production is fairly insensitive to changes in most viral entry, assembly, and release parameters, but highly sensitive to some viral transcription and translation parameters. Furthermore, we found a cooperative benefit to pairwise targeting of transcription and translation, predicting that combined targeting of these processes will be especially effective in inhibiting viral production.

## Main Text

The ongoing global pandemic caused by severe acute respiratory syndrome coronavirus 2 (SARS-CoV-2) has resulted in more than 4 million confirmed cases and 300,000 deaths worldwide (*1*). The most common symptoms of the illness caused by SARS-CoV-2, COVID-19, include fever, cough and fatigue (*2*). The clinical presentation can range from asymptomatic to fatal, with severe cases rapidly progressing to pneumonia, acute respiratory distress syndrome (ARDS), and organ failure (*3*). There is currently no vaccine for the disease, and development could take 12 to 18 months (*4*). Therefore, there exists a critical need for effective therapeutic interventions to minimize the transmission and severity of SARS-CoV-2.

Despite increased expenditures on research and development, less than one out of every ten therapeutic drugs that enter phase I clinical trials eventually gains FDA approval (*5*). To accelerate the development process for COVID-19, the initial focus has been on repurposing approved drugs and biologics. However, owing to the large number of possible therapeutics, and the need for rapid testing, there has not been extensive preclinical testing on SARS-CoV-2 specifically. While these therapeutics have been deployed in combinations of two, three, or four in more than 800 clinical trials globally (*6*), it is not clear which drugs, possibly in combination with others, would, in principle, be most effective. For other RNA viruses, such as HIV, single therapies have not been successful as a result of the virus’ ability to rapidly evolve and develop resistance to antivirals (*7*), thus driving the need for combination therapies (*8*). With more than 100 distinct agents currently in trials to treat COVID-19, even a two drug combination has over 10,000 possible combinations that could be tried, raising the question of how to rationally focus clinical trials on the single agents and combinations that are most likely to be effective.

Biophysical modeling has the potential to help rationally guide the development of therapeutic interventions for SARS-CoV-2 by identifying key model parameters for effective targeting. In addition, modeling can potentially be used to identify combination therapies, predict clinical trial outcomes, stratify patients, and identify potential source(s) of variable patient-to-patient outcomes. Here we present a biophysical model for the SARS-CoV-2 viral cycle and identify the single and combination parameters with the highest sensitivity, representing the ideal targets for therapeutic intervention to inhibit virus production.

The biophysical model of the SARS-CoV-2 viral life cycle was constructed based on SARS-CoV literature that included processes underlying viral entry, genome transcription, genome translation, virion assembly, and virion release (Fig. 1; reviewed in (*9, 10*)). Mass action and chemical rate equations were used to mathematically describe the system using an approach similar to that taken previously to model the life cycle of other viral systems (reviewed in (*11*); see Materials and Methods), and the resulting series of ordinary differential equations solved numerically. As an initial condition we assumed that a single virion was bound to the surface of an individual cell, which was then internalized to initiate the replication cycle. As data specific to the viral cycle to SARS-CoV-2 are limited, we primarily used experimental observations from SARS-CoV to inform model assumptions and parameterization (Table 1). We assume that it is reasonable to use SARS-CoV data based on the high degree of genomic similarity between the two viruses (*12–14*). Using the base parameter values, the model reproduces viral production on a timescale and of a quantity consistent with experimental observations (Fig. 2). Importantly, the model reproduces a number of experimental observations without parameter adjustment (Fig. 2B); for example, the model naturally predicts that ∼10% of the RNA will be negative sense (Fig. 2). Based on this, we conclude that the model provides a suitable tool to identify points of interest for therapeutic intervention, i.e. those parameters and the associated subprocesses that are particularly sensitive to perturbation.

**Table 1.**
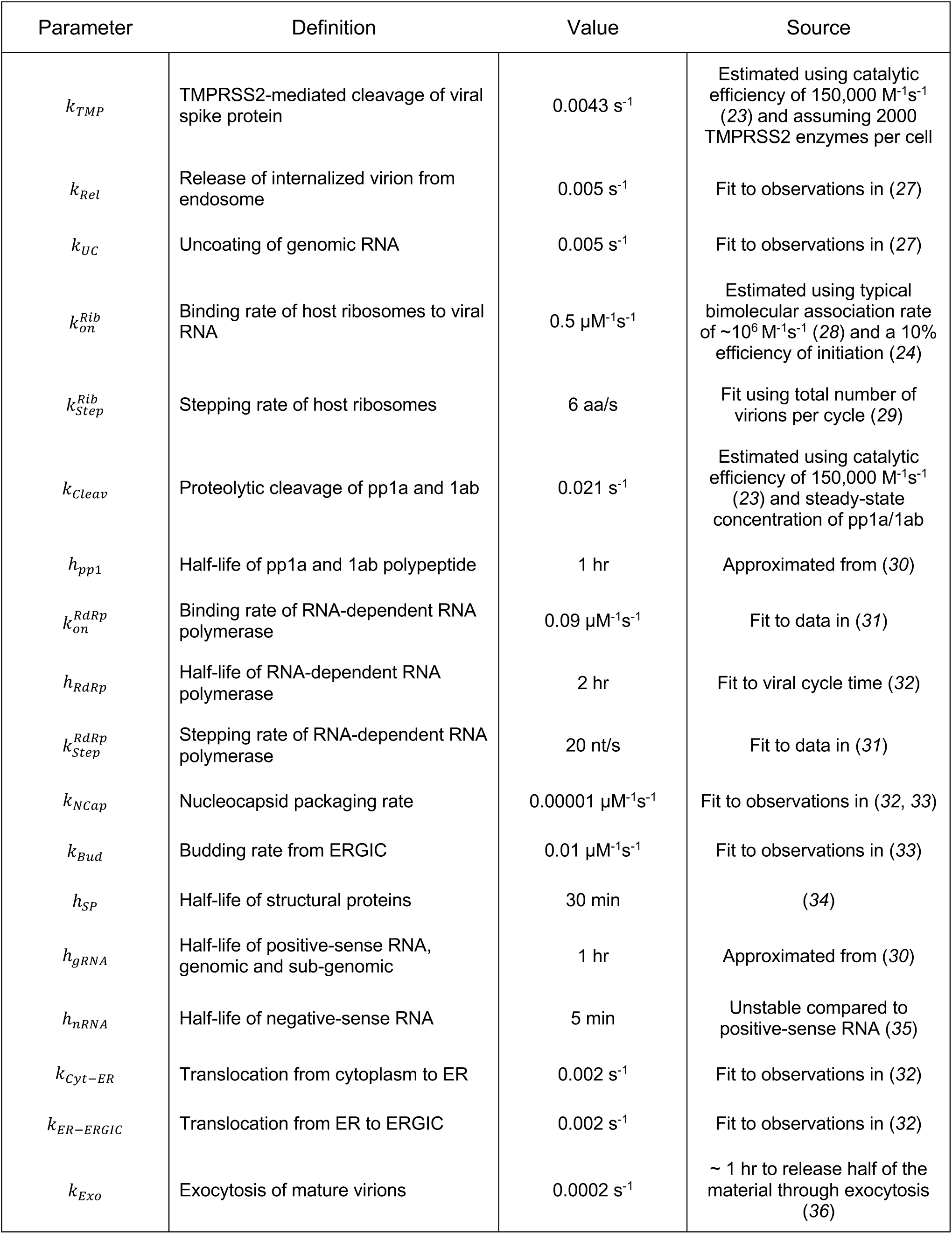
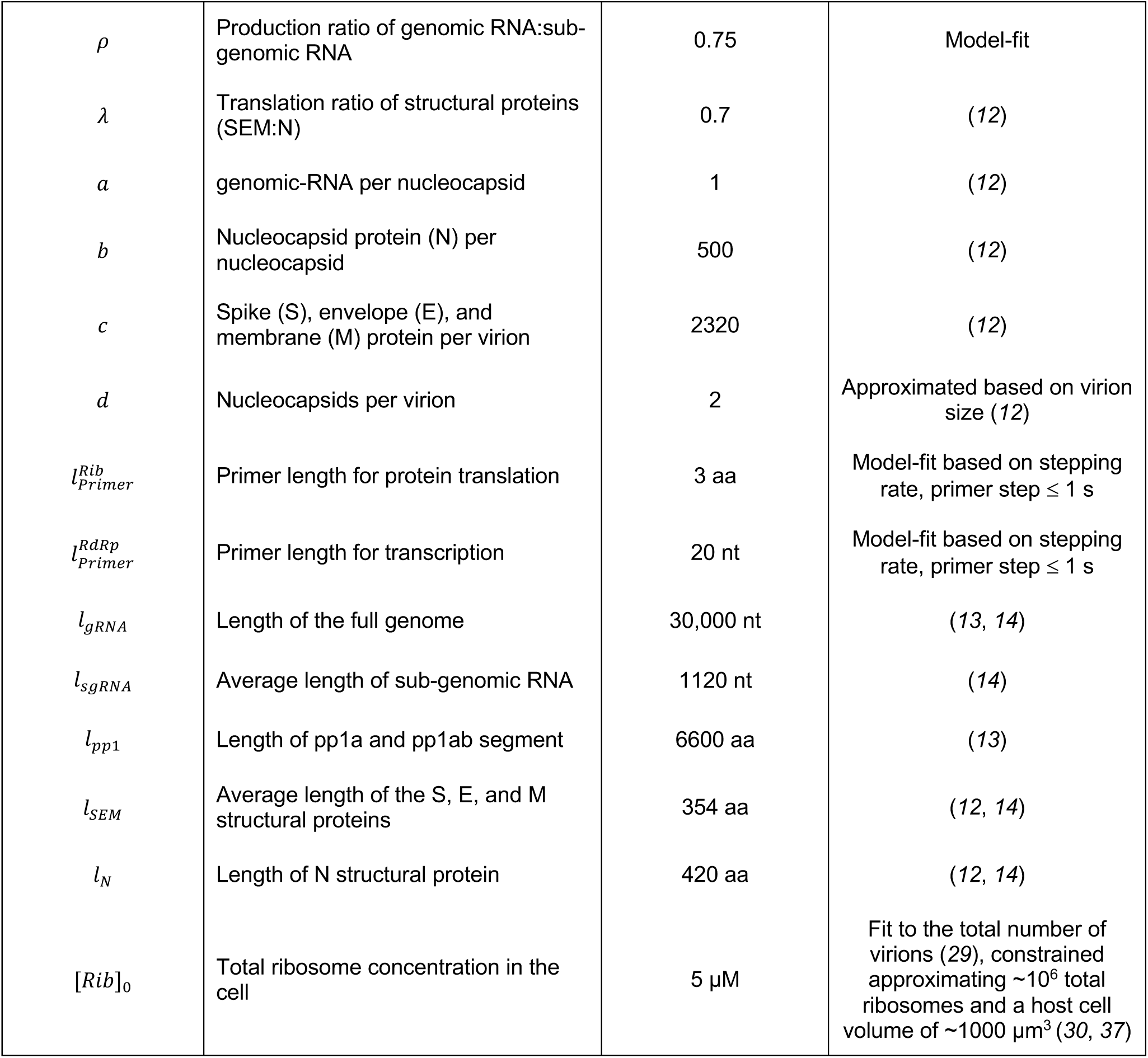
Model parameters.

**Figure 1.**
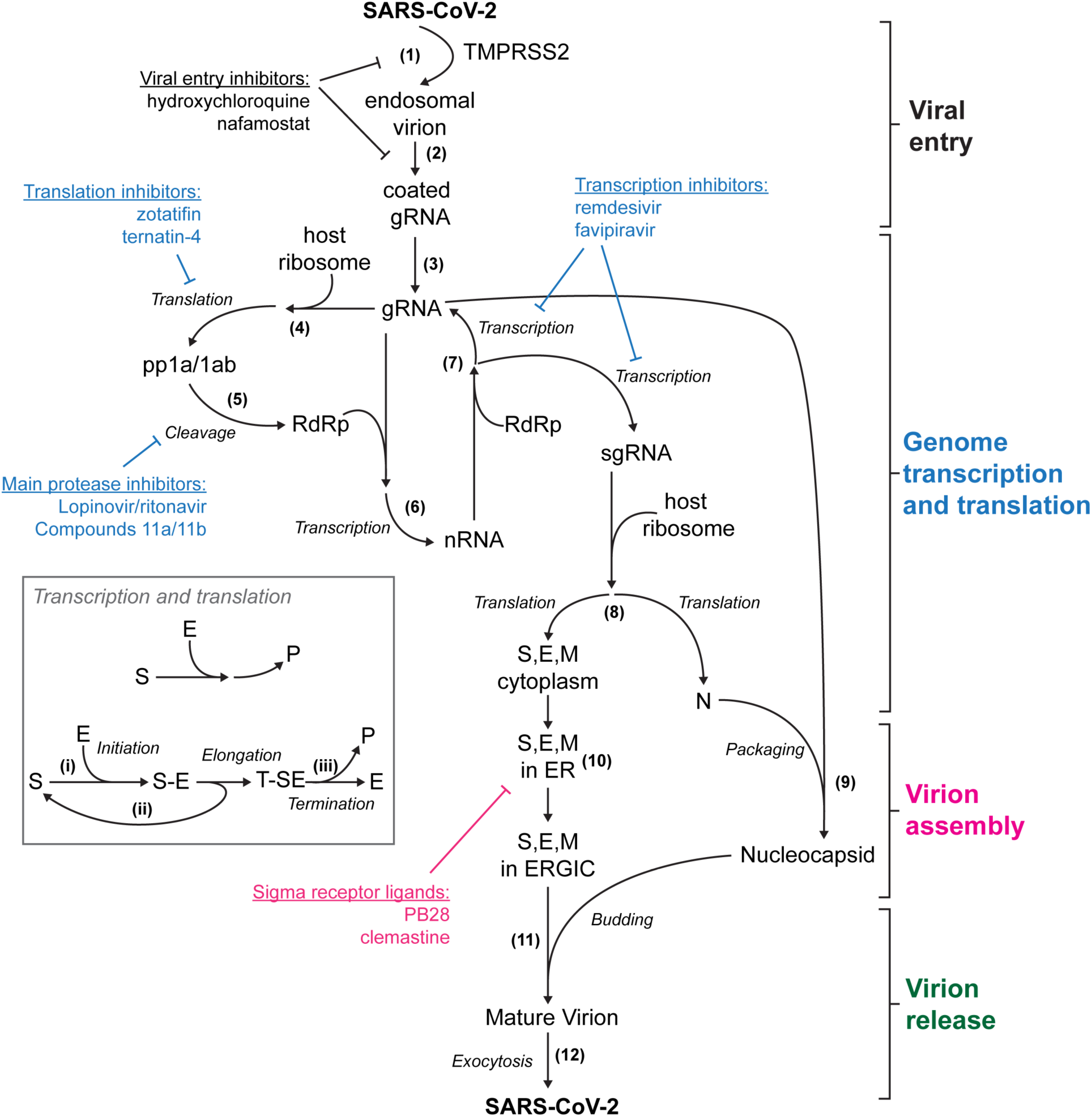
Model for the viral cycle of SARS-CoV-2. A single SARS-CoV-2 virion bound to the membrane of a target cell is internalized via TMPRSS2-mediated cleavage of the spike protein **(1)** followed by endosomal release **(2)** and uncoating **(3)** of the viral genome. Once the full-length genome is internalized, host ribosomes bind to open reading frames to translate the polypeptides pp1a and pp1ab **(4)**. Part of pp1a/1ab encodes the main protease (Mpro) involved in cleaving the polypeptides into the non-structural proteins that make up the replicase-transcriptase complex involved in replicating the virus, including the RNA-dependent RNA polymerase (RdRp) **(5). (6)** The RdRp first creates negative-sense RNA (nRNA) templates from the positive-sense genomic RNA (gRNA), which are then used to replicate the full-length gRNA as well as sub-genomic messenger RNAs (sgRNA) **(7)** encoding the structural proteins necessary to build the mature virion. **(8)** sgRNA is translated by the host ribosomes to make the structural spike (S), envelope (E), membrane (M), and nucleocapsid (N) proteins. **(9)** Multiple N proteins bind to the gRNA to package the nucleocapsids. **(10)** While initially translated in the cytoplasm, S, E, and M proteins are translocated through the endoplasmic-reticulum (ER) and ER-Golgi intermediate complex (ERGIC), where buds from the ERGIC eventually encapsulate nucleocapsids to form the mature virion **(11). (12)** Mature virions are then released into the surrounding tissue through exocytosis. Inset: detailed diagram of steps in transcription and translation. **(i)** Initially the RdRp or ribosome (enzyme) binds to the primer region of the respective RNA segment (substrate). **(ii)** Once the primer segment is completed, the substrate is released such that it is available for binding to the primer region by another enzyme. Meanwhile, the initial enzyme continues with elongation steps, polymerizing the reaction product. **(iii)** Upon complete polymerization of the product (RNA segment or protein), transcription/translation is terminated, and the enzyme and resulting product are released. Figure is annotated with example therapeutics to show their approximate point of influence on the viral cycle (*16, 22, 25, 26*). Examples are not exhaustive. Color of the text indicates the part of the viral cycle each example is associated with.

**Figure 2.**
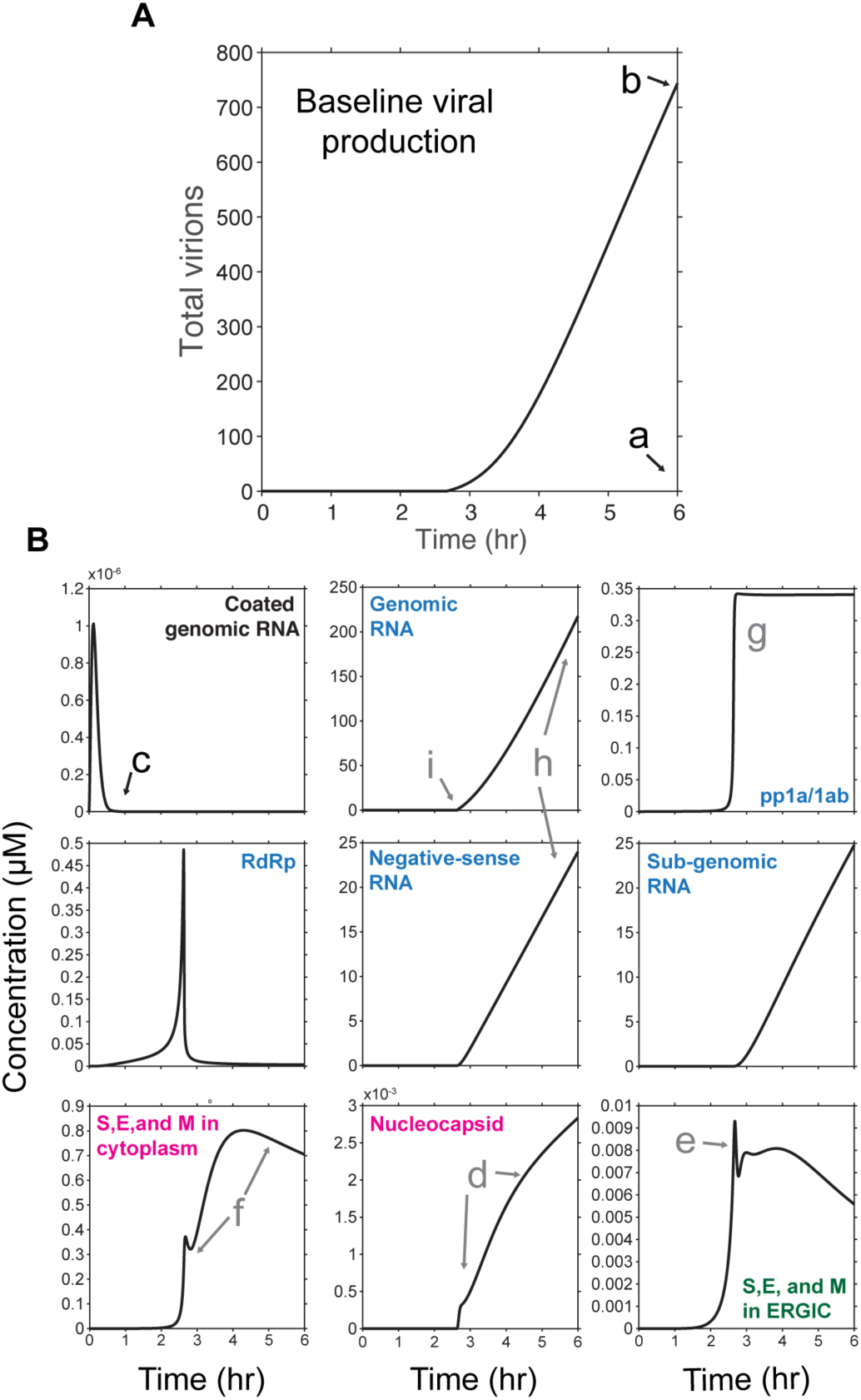
Baseline parameter values reproduce viral replication consistent with previous observations. A) Viral production as a function of time for the baseline parameters shown in Table 1. B) Concentration as a function of simulated time is shown for multiple species within the model. Specific species of interest are indicated within each subpanel. Color corresponds to the point of the viral cycle indicated in Figure 1. Letters indicate examples of model output that is consistent with experimental observations. Black text indicates those observations used to fit model parameters, while those in gray are observations that naturally occur as outputs of the model without parameter adjustment.

To assess these points of interest in the viral cycle, we performed a sensitivity analysis for each parameter by systematically increasing and decreasing their values from baseline by up to three orders of magnitude (1000-fold) while holding all other parameters constant (Fig. 3).

**Figure 3.**
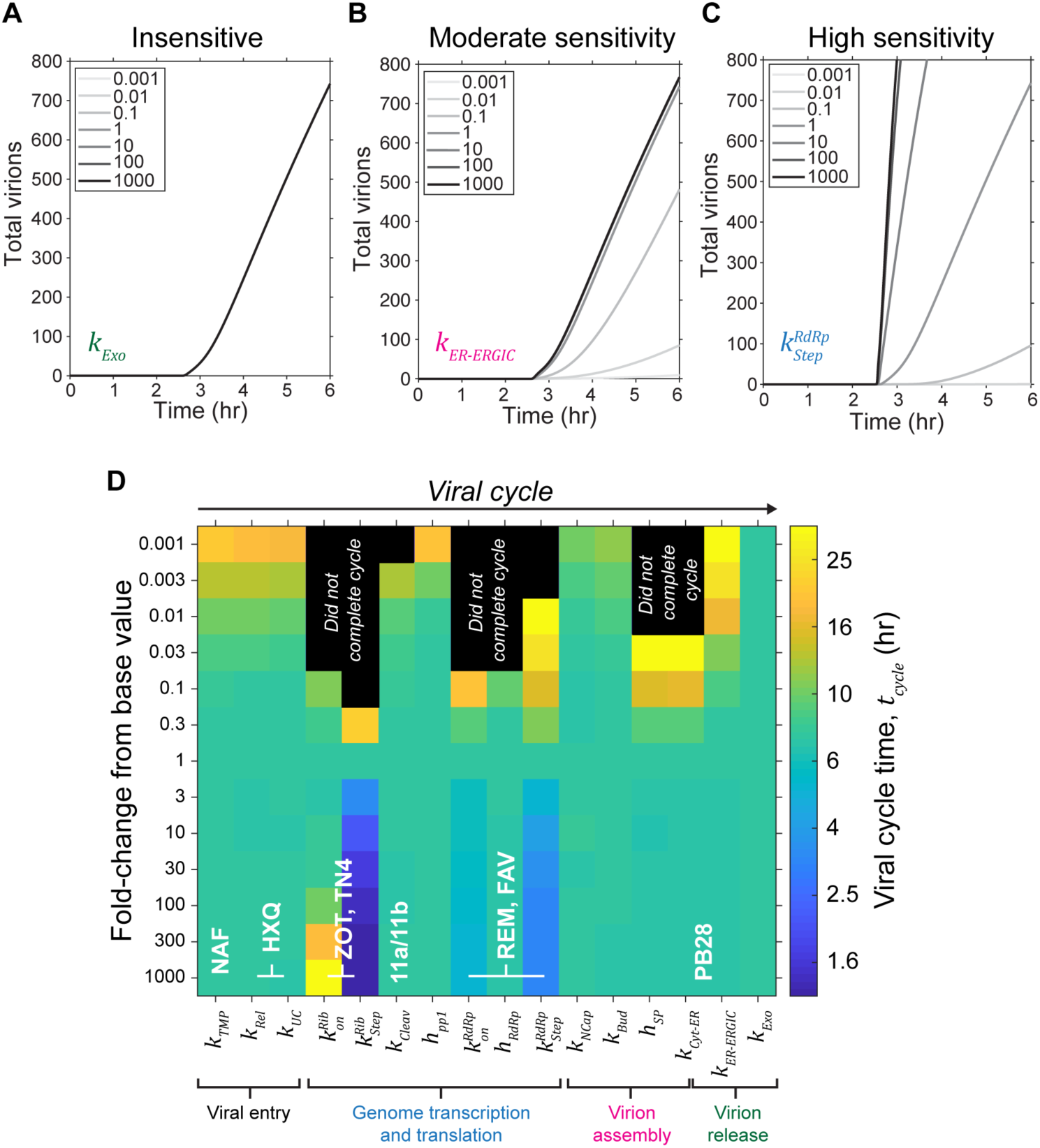
Sensitivity analysis reveals parameter targets of high interest. (A-C) Examples of insensitive (A), moderate sensitivity (B), and high sensitivity (C) parameters are shown. Inset: fold-change in parameter value relative to the baseline value. Color of the parameter text represents the point of the viral cycle indicated in Figure 1. D) Viral cycle time, estimated as the time to produce 1000 virions is shown as a function of varying parameter values. From left to right, parameters are sorted in the order in which they occur in the viral cycle. Black squares indicate parameter values where the model did not produce 1000 virions within 48 hrs. Figure is annotated with example therapeutics from Figure 1 to show their approximate point of influence on the model parameters. Examples are not exhaustive. NAF - nafamostat; HXQ - hydroxychloroquine; ZOT - zotatifin; TN4 - ternatin-4; 11a/11b - compounds 11a and 11b; REM - remdesivir; FAV - favipiravir.

Parameters specific to virion stoichiometry or RNA segment length were ignored in this analysis as they are likely difficult to target clinically. Using the viral cycle time *t*_*cycle*_, defined as the time to produce 1000 virions, as a readout of viral replication rate, we examined viral production as a function of individual parameter values. We found that parameter perturbation resulted in a variety of responses, ranging from insensitive to highly sensitive (Fig. 3A-C). As seen in Figure 3D, viral production was most sensitive to perturbation of many, but not all, parameters related to genome transcription and translation. While two parameters specific to viral packaging were able to eliminate viral production, the majority of parameters related to viral entry, packaging, and release were comparably insensitive to changes from the base values (Fig. 3D). While it is theoretically possible to inhibit viral production via targeting of the insensitive parameters, it is necessary to achieve a high level of inhibition compared to those that exhibit high sensitivity (Fig. S1). For example, parameters specific to viral entry and packaging required 100-1000x level of inhibition in order to influence viral production (Figs. 3D and S1A, C), while parameters specific to replication completely inhibited viral production after only a 10x effect (Figs. 3D and S1B). Based on these observations, we conclude that transcription and translation represent high sensitivity targets for therapeutic inhibition.

Due to the ability to rapidly evolve, viral diseases are often treated with combination therapies. While an exhaustive combinatorial analysis is difficult or impossible to conduct experimentally due to time and resource limitations, it can quickly be implemented *in silico*. To identify potential combinations that could cooperate to inhibit viral production in the model, we performed a pairwise sensitivity analysis of the model parameters. Parameter pairs were either coordinately increased or decreased from their base value and then scored based on the results (see Materials and Methods). To score parameter combinations we defined a sensitivity (*S*) and range (*R*) value (Fig. 4A), similar to our previous approach for another biophysical model (*15*). The sensitivity S is a measure of how viral production scales with changes to the base parameter values, while *R* is a measure of the magnitude of perturbation necessary to produce a maximal effect. Ideally, interventional therapies would have a strong effect (large *S*) with minimal perturbation (small *R*). Therefore, we scored parameter pairs by dividing the resulting *S* value by the value of *R* (*S/R*). As seen in Figures 4B and S2, several parameter combinations completely eliminated viral production with only a <10-fold effect. Similar to the single parameter sensitivity analysis, parameters related to transcription and translation once again emerged as the highest sensitivity targets; the highest scores were parameter combinations specific to the host ribosomes and the viral RNA-dependent RNA polymerase (RdRp). One other noteworthy pair that completely eliminated viral production was the half-life of the structural proteins (*h*_*SP*_), which comprise the mature virion, and the rate at which those proteins are translocated from the cytoplasm to the endoplasmic reticulum (*k*_*Cyto-ER*_). While the majority of high scoring combinations were simply the result of pairing with an already high sensitivity parameter 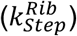, several combinations resulted in cooperating effects between the two parameters (Fig. 4C). As seen in Figure 4C, the greatest amount of cooperativity occurred when combining parameters specific to transcription and translation together. Based on these results, we conclude combination targeting of transcription and translation can lead to enhanced effects on viral production, beyond targeting each individually.

**Figure 4.**
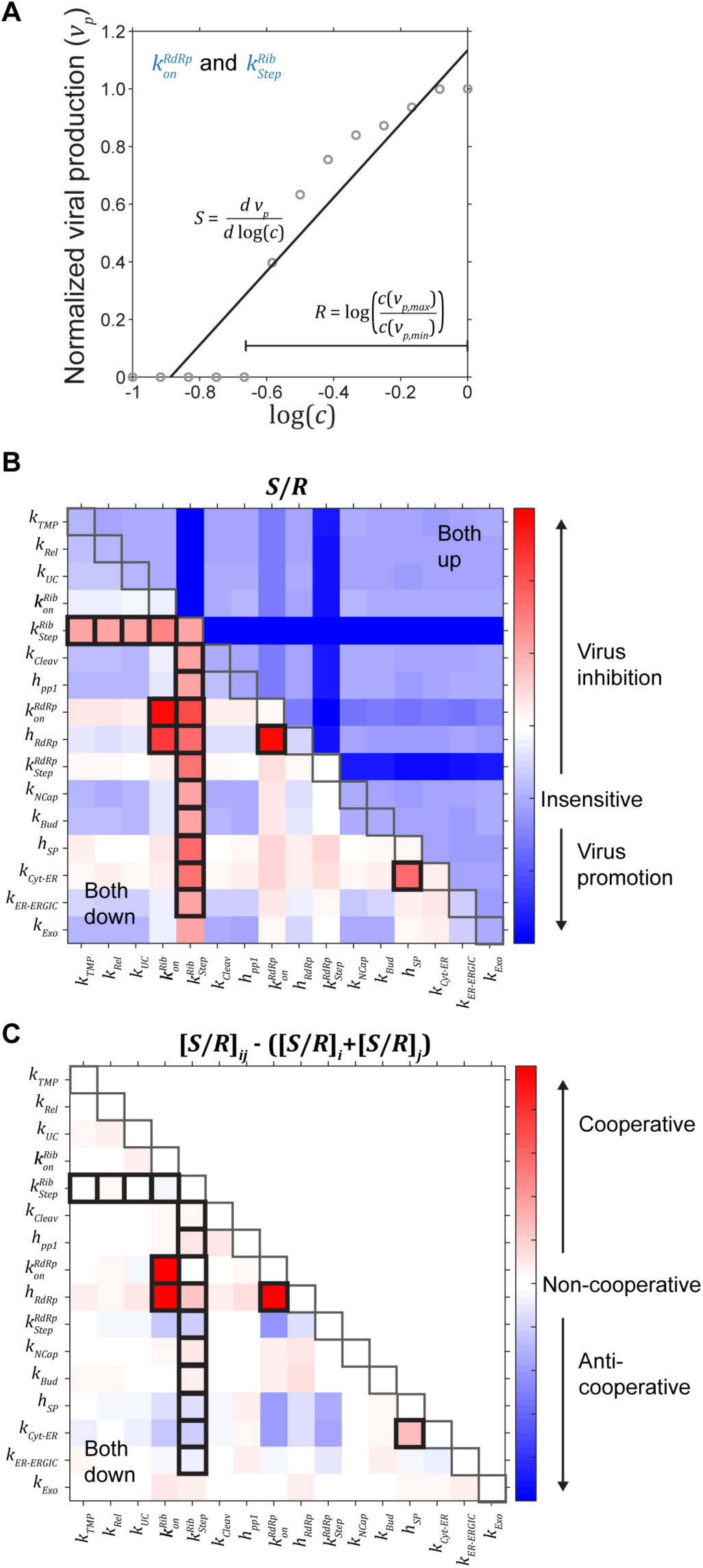
Pairwise sensitivity analysis reveals combinations of high interest. A) Example output used to score sensitivity (*S*) and range (*R*) is shown. *S* is estimated as the slope of the normalized viral production (*p*) versus the logarithm of the scaling parameter (*c*). As a result of the analysis approach, viral inhibition (as shown here) results in a positive slope, or *S* value, while promotion of viral production results in a negative *S* value. *R* is the difference between the scaling parameter values where maximum and minimum viral production occur. B) Pairwise scoring of sensitivity and range (*S*/*R*). Gray boxes indicate the diagonal where only a single parameter is scaled. Relative to the diagonal, the upper-right region is where both parameter values were increased, while the lower-left region is where both parameter values were decreased. Bold boxes indicate parameter combinations that eliminated viral production (*p*_*min*_ = 0). C) Cooperative parameter effects are estimated by the difference between the combined effect ([*S*/*R*]_*ij*_) and the sum of the individual effects ([*S*/*R*]_*i*_ + [*S*/*R*]_*j*_). Since increasing parameter values (upper-right) did not result in viral inhibition (B), this region was omitted. Positive values (red) resulted when the combined effect was stronger than the sum of the individual effects. Negative values (blue) resulted when the combined effect was weaker than the individual effects. Gray and bold boxes are the same as in B.

Overall, our modeling of the SARS-CoV-2 life cycle, parameterized using published SARS-CoV literature, shows that theoretically there are opportunities for therapeutic interventions that significantly inhibit the viral cycle. In particular, the sensitivity analysis identified several parameters in the middle of the viral cycle, specific to genome transcription and translation, that present the best opportunities for inhibiting viral production. By comparison, parameters specific to viral entry, virion assembly, and virion release were less sensitive and therefore are less promising as targets for inhibiting viral production. The model further identifies potential combination targets that would cooperatively inhibit viral production. For example, the combined effects of targeting both the stepping rate of the host ribosome, 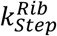, and the binding rate of RdRp,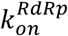, may halt the viral cycle even with modest 10X effects. Such a pairwise analysis would be difficult to exhaustively test experimentally, especially in the clinic, since there are 16^2^ = 256 possible combinations.

In addition to identifying novel target opportunities within the viral cycle, biophysical modeling may also provide insight into current and emerging therapeutic approaches. Several existing antiviral drugs are being evaluated for their efficacy in treating COVID-19 (*16*). It is interesting to note that remdesivir, which was recently approved for emergency use in the U.S. and Japan, acts to disrupt genome transcription by interfering with the viral RdRp (*17, 18*), and thus is one example of a therapeutic intervention that interferes with an area of high sensitivity identified by the model. By contrast, hydroxychloroquine, which inhibits viral entry by increasing endosomal pH and affecting terminal glycosylation of the ACE2 receptor (*19*), is predicted by the model to be less likely to be effective since drugs that act on viral entry would require exceptionally high suppression to achieve appreciable effects on viral production. Viral entry inhibitors have been used to treat HIV (*20*) and influenza (*21*), therefore it may yet be feasible to target viral entry. Additionally, we note that therapeutics predicted to be less effective in the viral cycle model, may have potent effects elsewhere in disease progression, e.g. in the immune response. Thus, ineffectiveness in inhibiting viral production does not preclude therapeutic effectiveness overall.

Finally, the recent host-virus protein interaction map reported by Gordon et al. (*22*) identified two classes of SARS-CoV-2 targeting opportunities: 1) inhibitors of protein translation and 2) regulators of Sigma1 and Sigma2 receptors. Based on our modeling, we predict that therapeutic targeting of the Sigma1 and 2 receptors is not likely to be effective as it would presumably interfere with the virion assembly and release steps, which are relatively insensitive (Fig. 3D). By contrast, we predict that targeting protein translation is more likely to be effective due to the high sensitivity of viral cycle time to translation-associated parameters (Fig. 3D). Everything else being equal, the model predicts that transcription inhibition combined with translation inhibition would be an especially effective combination (Fig. 4B-C). Altogether, our model provides a framework for understanding viral cycle dynamics and identifying the therapeutic opportunities that are most likely to be effective in inhibiting viral production.

## Acknowledgements

The authors thank David Largaespada and Jonathan Sachs for helpful discussions and thank Ismail Guler for technical assistance.

## Author contributions

All authors contributed to the development and the design of the model. B.T.C. wrote the computer code and performed all analyses. B.T.C, C.D., and D.J.O wrote the manuscript. All authors commented on and contributed to the manuscript.

## Data and materials availability

All computer code developed in this manuscript will be made available on the Odde Lab webpage at http://oddelab.umn.edu/software.html.

## Materials and Methods

The system of ODEs describing the viral cycle was solved using the *ode15s* solver in Matlab (R2019a; The Mathworks, Natick, MA).

### Viral entry

Upon initialization the model is assumed to have a single viral particle (SARS-CoV-2) bound to angiotensin-converting enzyme 2 (ACE2) on the cell membrane (SCoV2-ACE2). SARS-CoV-2 is internalized into endosomes as the spike protein is cleaved by transmembrane protease, serine 2 (TMPRSS2) at a first order-rate (*k*_*TMP*_).

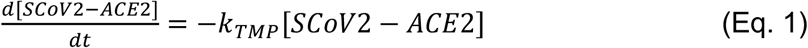

This reaction and other proteolytic reactions in the model are simplified to the product of a first-order rate constant *k* and the concentration of the enzyme ([*E*]) where *k =* (*k*_*cat*_*/K*_*M*_)[*E*]_SS_. This assumption is true when the substrate concentration is much less than the Michaelis-Menten constant value, [*S*] << *K*_*M*_. For each reaction we assumed a catalytic efficiency (*k*_*cat*_*/K*_*M*_) equal to 150,000 M^-1^s^-1^, the median value for a range of enzymes (*23*). The steady-state concentration of the enzyme, [*E*]_*SS*_, was estimated using the baseline parameter values in Table 1. Endosomal SARS-CoV-2 is then released and the genomic RNA uncoated with the first-order rates *k*_*Rel*_ and *k*_*UC*_, respectively.

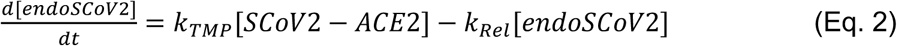

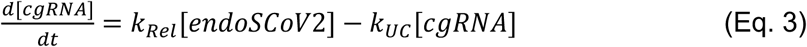

### Transcription and translation

After the full-length genomic RNA (gRNA) is released and uncoated, host ribosomes bind to open reading frames to begin translation of pp1a and pp1ab. In each instance, transcription and translation is modeled as a multistep process (Figure 1 of the main text; inset), similar to that outlined in (*24*).

i. Initial binding of the enzyme (Ribosome or RNA-dependent RNA polymerase) to its respective substrate (RNA) is assumed to be irreversible and is the product of the second-order association rate (*k*_*on*_) and the concentrations of the enzyme and its substrate.
ii. The enzyme completes the primer sequence at rate *k*_*Prime*_, which is determined by the stepping rate of the respective enzyme (*k*_*Step*_) and the length of the primer sequence (*l*); *k*_*Prime*_ *= k*_*Step*_*/l*. This first step is fast (∼0.5-1 s), and once completed the substrate is released such that multiple enzymes can bind to a single strand of RNA. The actively polymerizing translation or transcription complex (T-SE) then continues through the elongation process.
iii. For each active T-SE, the rate of termination (*k*_*Term*_) is proportional to the length (*l*) of the sequence and the stepping rate; *k*_*Term*_ *= k*_*Step*_*/l*. Upon termination, the enzyme unbinds from the substrate and the product is released.

For the specific example of translating pp1a/1ab by the host ribosomes, this multistep process results in the following equations

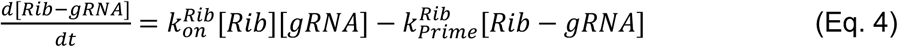

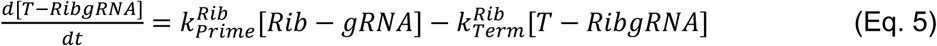

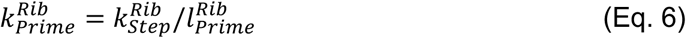

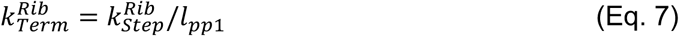

Additionally, the polypeptides pp1a/1ab are then cleaved at first-order rate *k*_*Cleav*_ to release the non-structural proteins making up the replicase-transcriptase complex, specifically the RNA-dependent RNA polymerase (RdRp), and is degraded with a half-life of *h*_*pp1*_.

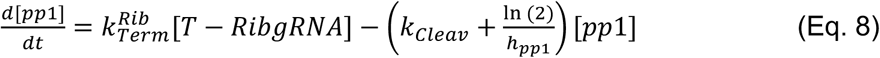

The multistep process outlined above repeats for production of negative-sense RNA (nRNA) templates,

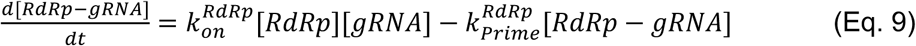

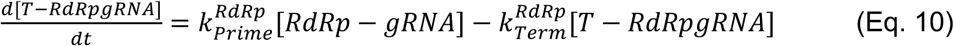

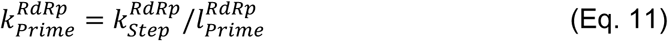

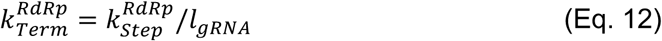

The resulting nRNA templates are then used to transcribe the full-length gRNA and sub-genomic RNA (sgRNA) encoding the structural proteins. Additionally, nRNA is degraded with a half-life of *h*_*nRNA*_

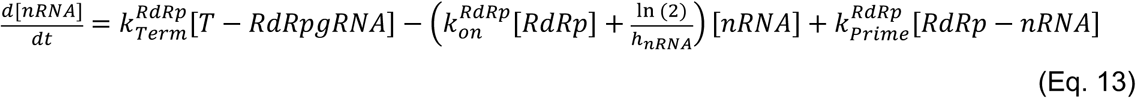

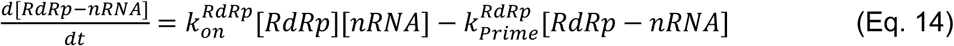

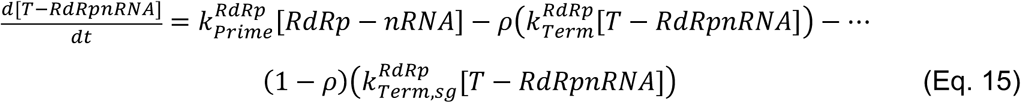

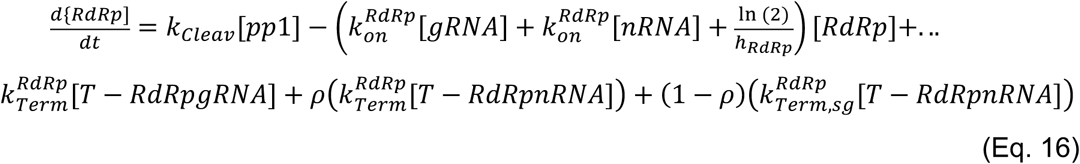

Here, *ρ* is the fraction of transcription complexes producing full-length gRNA as opposed to sgRNA. sgRNA is produced at a faster rate due to the shorter sequence length such that

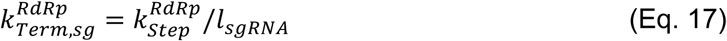

Where *l*_*sgRNA*_ is the average length of sgRNA coding the structural proteins, which is weighted by the relative length and stoichiometry of each protein. This then completes the cycle for replicating the full-length gRNA. An additional term is added for the coating of the genome with nucleocapsid (N) protein in preparation for encapsulation in the mature virion, where *k*_*NCap*_ is the second-order rate of producing nucleocapsids.

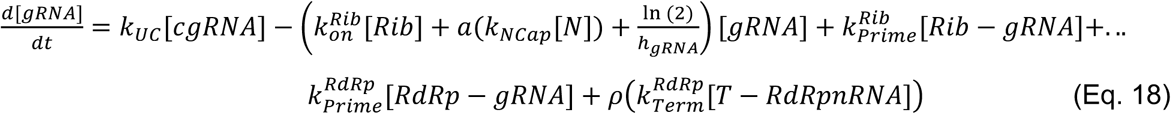

The final multistep process is the translation of sgRNA by host ribosomes to produce the structural proteins making up the mature virion, the spike (S), envelope (E), membrane (M), and nucleocapsid protein (N). Additionally, sgRNA is degraded with the same half-life as gRNA (*h*_*gRNA*_).

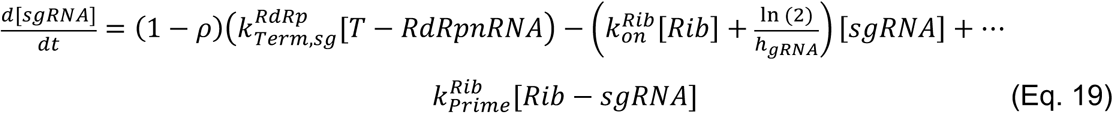

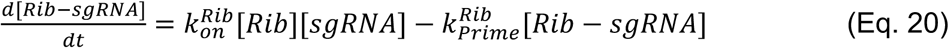

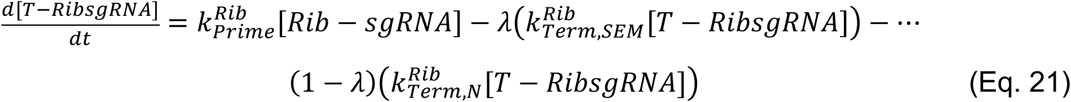

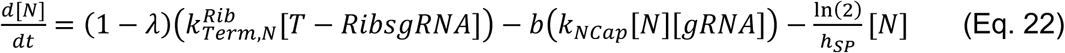

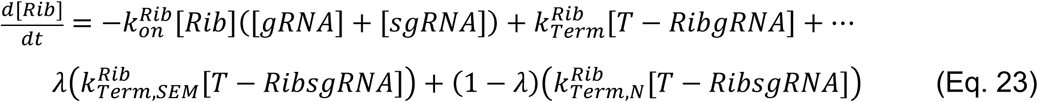

where

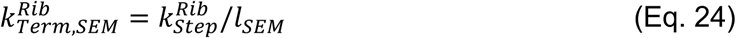

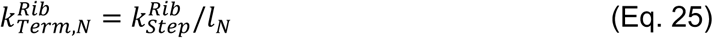

In the above equations, *λ* is the ratio of producing S, E, or M proteins relative to N protein. The value of *λ* is determined by the stoichiometries of each. Furthermore, the translation rate of S, E, or M protein is assumed to be proportional to the final stoichiometry in the mature virion, and therefore they can be lumped together, simplifying the handling equations.

### Virion assembly and release

S, E, and M proteins are polymerized in the cytoplasm but are then translocated through the endoplasmic reticulum (ER) and ER-golgi intermediate complex (ERGIC) at first-order rates *k*_*Cyt-ER*_ and *k*_*ER-ERGIC*_, respectively.

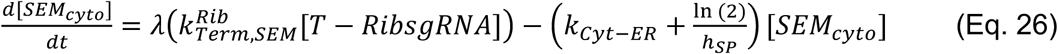

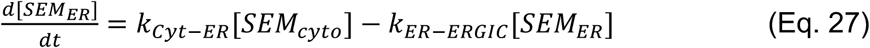

Budding from the ERGIC at rate *k*_*Bud*_ forms the mature virion from the encapsulation of nucleocapsids. Mature virions are then released via exocytosis at first-order rate *k*_*Exo*_.

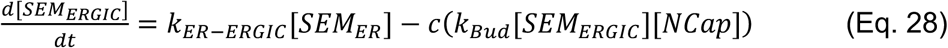

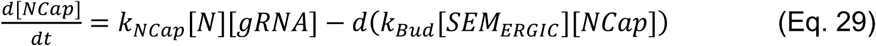

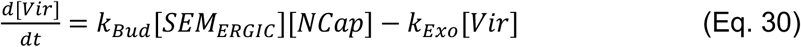

### Sensitivity analysis

To initially assess model sensitivity, we used the viral cycle time *t*_*cycle*_, defined as the time to produce 1000 virions, as a measure of viral production; the longer the cycle time, the lower the viral production. For single parameter changes (Figure 3 in the main text), individual parameters were adjusted via multiplication by a scaling constant (*c*), which was assigned values between 0.001 and 1000 on a logarithmic scale.

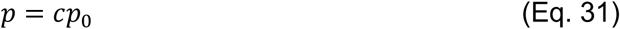

where *p*_0_ is the baseline value of the specific parameter.

Pairwise sensitivity analysis was performed using a similar approach to that described in (*15*), where a sensitivity (*S*) and range (*R*) value were assigned based on how model output scales with changes in parameter values. To perform the pairwise analysis, we defined viral production level by

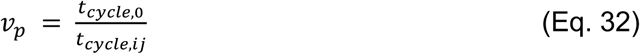

where *t*_*cycle*,0_ is the baseline cycle time and *t*_*cycle,ij*_ is the cycle time resulting after perturbation of the paired parameters. Parameter combinations were scaled using a constant (*c*), which was assigned values between 0.1 and 1 on a logarithmic scale. Both parameters were scaled down by multiplication with the constant

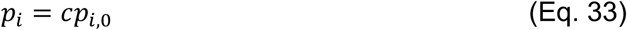

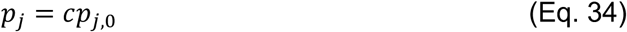

or scaled up by dividing by the same constant value.

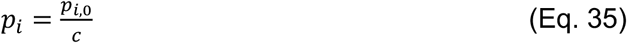

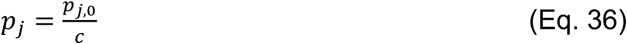

A sensitivity value was assigned by measuring the change in viral production as a function of the change in the parameter values.

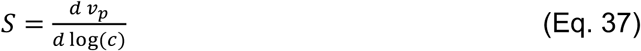

Thus, *S* is the slope of the plot of *v*_*p*_ versus log(*c*) (Figure 5A in the main text). Assigning values using Eqs. S30 and S35 resulted in positive values for parameters that inhibited viral production, increasing cycle time, and negative values for parameter changes that promoted viral production, decreasing cycle time. The *R* values were defined as the scale value between the maximum and minimum viral production level.

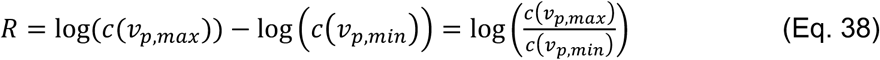

A minimum of a 10% change in viral production was required to assign a value, such that if *v*_*p,max*_ *– v*_*p,min*_ ≤ 0.1, then *R =* 1. We scored pairwise combinations by multiplying the sensitivity value by the reciprocal of the range value.

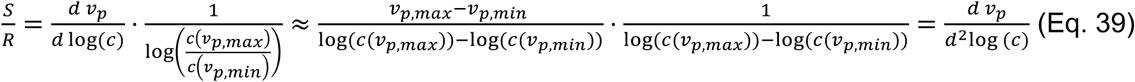

Doing so assured higher scores were assigned to those combinations exhibiting high sensitivity over a small range of values, the ideal characteristic for targets of therapeutic intervention. Individually, some parameters are insensitive while others are highly sensitive (Figure 3A-C in the main text). Therefore, we sought to assess whether high-scoring pairwise combinations were simply the combined effects of two sensitive parameters or if there was some amount of cooperativity occurring when both were targeted. We approximated parameter cooperativity as the difference between the pairwise score and the sum of the individual scores according to

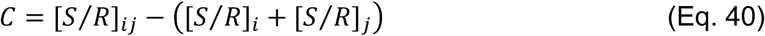

Thus, cooperative pairs will have a positive value according to Eq. 40, while those that are anti-cooperative will have a negative value.

### Modeling small molecule inhibitors

To simulate the effects of a small molecule inhibitor on individual parameters we scaled individual parameters (*p*) according to a Hill-function

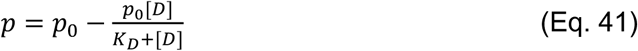

where *p*_0_ is the baseline value, [*D*]is the concentration of the theoretical small molecule inhibitor, and *K*_*D*_ is the dissociation constant, or the affinity, of the inhibitor for its target parameter. Simulated values for *K*_*D*_ are indicated in Figure 4 of the main text. Inhibitor concentrations were varied between 0.01 and 100 μM on a logarithmic scale. For each case, viral production was estimated according to Eq. 32 above. Levels of inhibition at each concentration come from the Hill-function in Eq. 41 and are equal to *p*/*p*_0_ for a given inhibitor concentration and *K*_*D*_ value.

