## Supplemental Information for "Biophysical modeling of the SARS-CoV-2 viral cycle reveals ideal antiviral targets"

Affiliations:
[1]Department of Biomedical Engineering, University of Minnesota, Minneapolis, MN 554455
[2]Department of Medicine, Division of Infectious Disease and International Medicine, University of Minnesota, Minneapolis, MN 55455
[3]Department of Surgery, Institute for Health Informatics, University of Minnesota, Minneapolis, MN 55455
[4]Department of Microbiology and Immunology, Center for Immunology, University of Minnesota, Minneapolis, Minnesota, 55414

Correspondence to:
[*], (612) 626-9980

**This PDF file includes:**

**Figures S1 to S2**

Supplementary Figures:

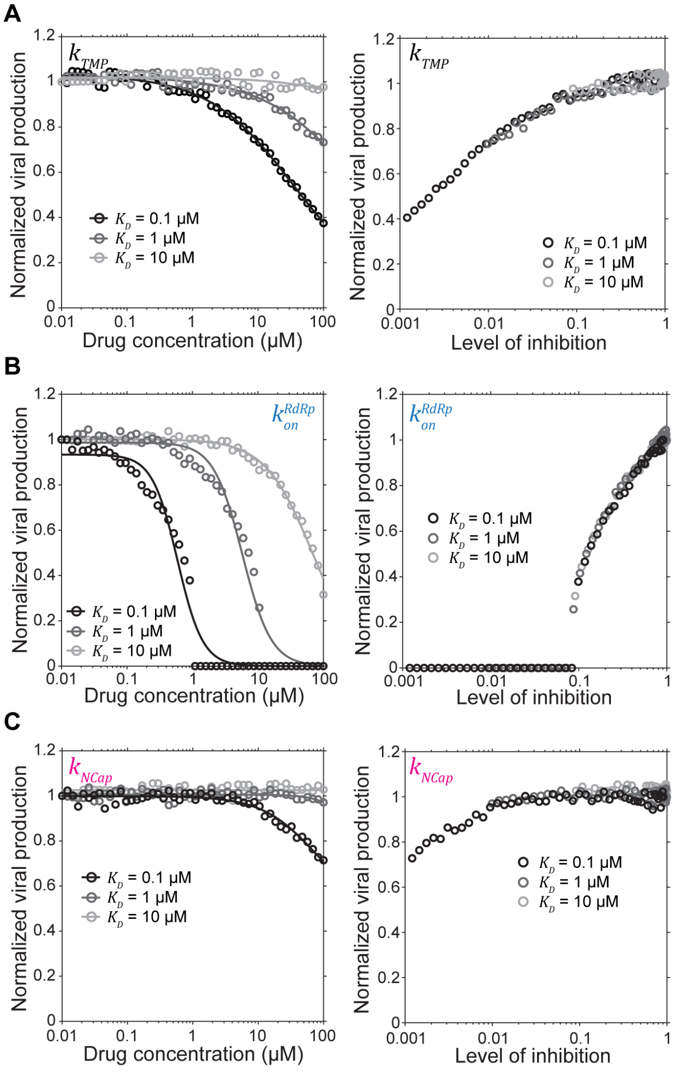


**Figure S1 - High sensitivity parameters represent ideal targets for small molecule inhibitors.** A-C) Simulated drug effects for theoretical small molecule inhibitors that target individual parameters. Left: viral production as a function of simulated drug concentration. Viral production was estimated by apparent cycle time, the time taken to reach 1000 virions, and was normalized to the cycle time in the absence of any drug effects. Lines are best-fit Hill-function. Right: viral production as a function of the level of parameter inhibition. $K_D$ values indicate the simulated affinity of the theoretical drug for its target parameter. Color of parameter text represents the point in the viral cycle corresponding to Figure 1 of the main text.

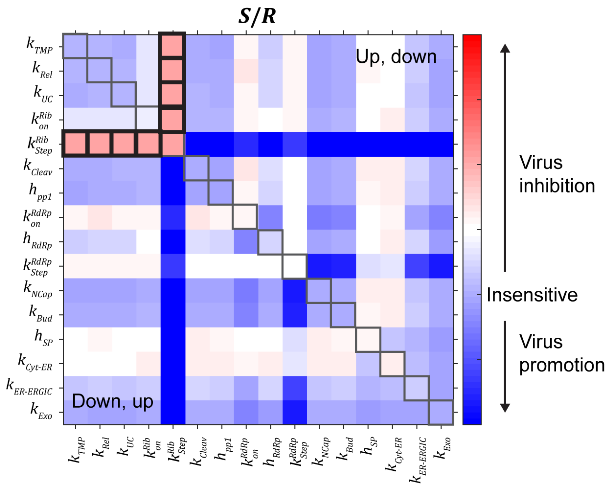


**Figure S2 - Extended pairwise sensitivity analysis.** Pairwise scoring of sensitivity and range ($S\,/R$) as performed in Figure 4 of the main text, however, here paired parameters were scaled in opposite directions. Gray boxes indicate the diagonal where only a single parameter is scaled. Relative to the diagonal, the upper-right region is where parameters on the y-axis were scaled up while those on the x-axis were scaled down. In the lower-left region, parameters on the y-axis were scaled down and those on the x-axis were scaled up. Bold boxes indicate parameter combinations that eliminated viral production ($p_{min}$ = $0$).
